# Birth weight and in-feed antibiotics alter the assembly and succession of the swine gut microbiota

**DOI:** 10.1101/2024.01.17.576096

**Authors:** Wenxuan Dong, Paul Oladele, Ruth Eunice Centeno-Martinez, Tessa Sheets, Brian Richert, Timothy A. Johnson

## Abstract

Understanding the principles of gut microbiota assembly and succession during host development is critical for effective gut microbiome manipulation to optimize host health and growth. The objective of this study was to conduct a high-frequency sampling of the swine gut microbiota from controlled groups of pigs to increase understanding of the dynamics of microbial community development. Here, a total of 924 fecal samples from 44 piglets (22 low-birth-weight, LBW; 22 normal-birth-weight, NBW) over 21 time points (1-41 days of age) collected every two days. Community composition, assembly, and succession was determined using 16S rRNA gene amplicon sequencing. Alpha diversity continuously increased during the suckling stage, yet no significant increase was observed during the days post-weaning. Post-weaning in-feed antibiotics consistently decreased microbial diversity and changed the community structure in both LBW and normal birth weight (NBW) piglets. Delayed post-weaning gut microbial community maturation was observed in LBW piglets compared with NBW. Heterogeneity of the gut microbial community between piglets linearly decreased over time, as revealed by the within-time Bray-Curtis dissimilarities. Individuality analysis on the relatively stable stage revealed that the gut microbiota composition of some individuals over time, and the abundance of most genera between individuals, were highly variable. Dirichlet multinomial mixtures analysis supported an age-dependent microbiota developmental pattern and identified the age-discriminatory taxa. The importance of stochastic processes in microbial community assembly increased over time within primary and secondary successions, despite the fact that the most dominant factors influencing community assembly were homogeneous selection and dispersal limitation, which are deterministic.

**IMPORTANCE:** Our understanding of the assembly and succession of the swine gut microbial community is limited, and scientific advancement in this interdisciplinary topic is hampered by individuality and transient dynamics. The solution to the above foundational questions is not only ecologically relevant but also useful for practical swine production. Our study addresses ecological processes shaping the swine gut microbiota between piglets with contrasting birth weights and receiving post-weaning antibiotics. Persistent gut microbiota immaturity in LBW piglets suggests that efforts to accelerate microbial community succession might improve LBW piglet growth performance and disease resistance. Intra-individual variance both in community structure and genus abundance during the post-weaning period indicates the importance of repeated measurements for reliable observations. Additionally, neutral (stochastic) processes increased as a factor of community assembly within each stage of pig growth, indicating that early intervention and multiple follow ups may be critical in manipulating the gut microbiota development.

## INTRODUCTION

The gut microbiota plays a critical role in food animal health and gut microbial variations could partially account for variations in feed efficiency, post-weaning illness, and growth performance (1–6). For these reasons, the importance of the gut microbiota is receiving increased attention in commercial swine production. Antibiotics, targeting the gut microbiota directly, have been widely used in animal husbandry for efficient pathogen control and growth promotion (7–9). However, antibiotic resistance has been considered as a global health problem; thus, limiting antibiotic use in food animals has increased in recent years (10–12). Although probiotics and/or prebiotics are promising approaches for modulating the gut microbiome to reduce the use of antibiotics in swine production, they cannot match the efficacy of antibiotics (13–16). Thus, a detailed understanding of gut microbiome development, especially under the antibiotics context, is critical to the development and successful use of probiotics and prebiotics as antibiotic alternatives. Low birth weight (LBW) piglets, with significantly reduced growth performance and increased disease susceptibility compared with their normal birth weight (NBW) peers, are benefitted by in-feed antibiotics even more than NBW piglets and provide a potential to explore the mechanism by which antibiotics promote pig health and growth (17–19). Different gut microbial communities, metabolism, and gene functions were observed in piglets with contrasting birth weights, confirming the correlation between gut microbiota and animal phenotypes (20, 21). However, little is known about the gut microbial community assembly and succession in LBW or antibiotic-treated piglets since previous studies mostly focused on normal birth weight and healthy pigs (22–24).

Understanding the gut microbial community assembly and succession is a fundamental ecological question and has a crucial role in the successful intervention of the gut microbiota. Primary succession begins postnatal with the arrival of pioneer microbes which shape the subsequent community assembly, and secondary succession happens postweaning with abrupt changes in the gut environment caused by a shift from liquid milk to solid feed (25, 26). Longitudinal studies have determined the microbiome development of both humans and pigs and have provided a foundation for mechanistic investigations of microbes and host health (27–30). However, a clearer view of the development of the human gut microbiome was revealed using fecal samples obtained on a near-daily basis (31, 32). High-frequency sampling with shorter time intervals is critical in determining the continual dynamics of the gut microbiome by capturing both the individuality and convergence of the community between individuals rather than “snapshot” samples (33, 34). Swine gut microbiome research also requires a detailed description of microbial succession in the gut to serve as a benchmark to future studies.

In this study, we collected fecal samples every two days from 44 piglets, beginning at the neonatal stage until three weeks postweaning (21 timepoints). The fecal microbiota were assessed using 16S rRNA gene sequencing to determine how birth weight and postweaning in-feed antibiotics affected the microbiota over this growth-period of piglets. Given that in humans, persistent gut microbiota immaturity has been observed in malnourished children (35, 36), we hypothesized that gut microbiota development is slower in LBW piglets than NBW and in control compared to piglets given in-feed antibiotics. We found an age-dependent microbiome assembly and successional pattern and persistent alterations by antibiotics. We also quantified the microbiota maturation curve using microbiota age, and the relative importance of ecological processes underlying microbial community assembly.

## RESULTS

### Growth performance of piglets

Piglets with low birth weight (LBW) had a lower average daily gain (ADG) than normal birth weight (NBW) piglets during the suckling period (P < 0.001, Fig. 1A). Despite all weaning piglets having free access to feed and water, the growth gap between LBW and NBW piglets further widened after weaning, indicating no post-weaning compensatory growth in LBW piglets (Fig. 1B). In-feed antibiotics significantly increased the post-weaning ADG of both LBW and NBW piglets. Furthermore, weaning piglets fed antibiotic-containing diets had a lower incidence of diarrhea than the control group (Fig. S1).

**FIG 1.**
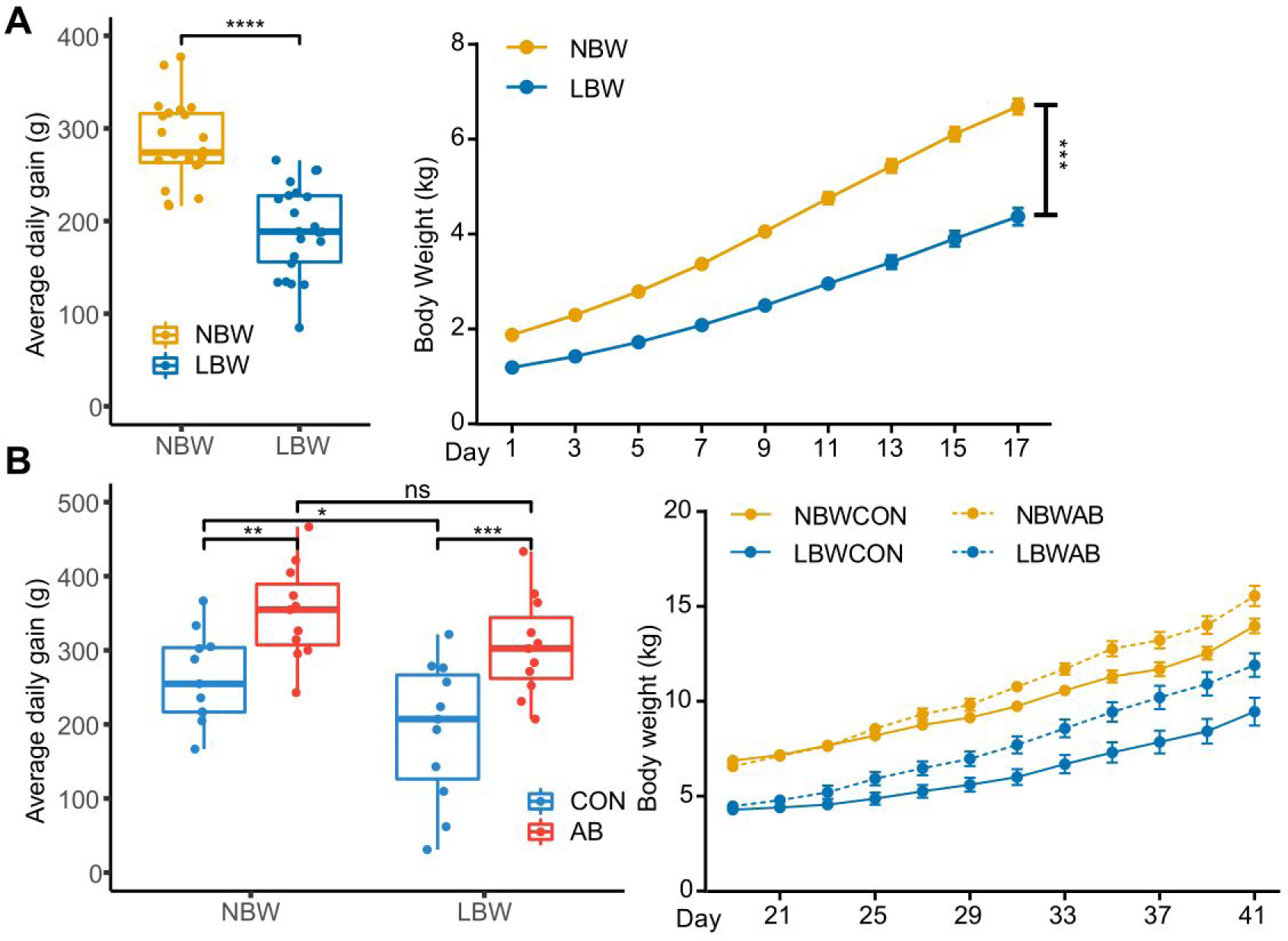
Average daily gain of piglets in suckling stage (A) and weaning stage (B). LBW, low-birth-weight; NBW, normal-birth-weight; AB, antibiotics; CON, control. *, P < 0.05; **, P < 0.01; ***, P < 0.001; ****, P < 0.001.

### Alpha diversity and taxonomic composition

A positive correlation between age and alpha diversity was observed in suckling piglets. Faith’s phylogenetic diversity increased over time in both LBW and NBW suckling piglets (Spearman’s rho = 0.70, P < 0.001, Fig. 2A). However, there was no linear correlation between age and alpha diversity after weaning (Fig. 2B). There were no significant effects of birth weight on Faith’s phylogenetic diversity in suckling piglets (Table S2). We used linear mixed-effects model including random intercepts and within-subject factor slopes for individual piglets nested by litter, and evaluated birth weight, age, and antibiotic treatment as fixed factors and gender as a cofactor. There were significant interaction effects between age and birth weight (P < 0.001), and between age and antibiotics use (P < 0.001) on Faith’s phylogenetic diversity in weaning piglets. After running post-hoc pairwise comparisons, we found that in-feed antibiotics significantly (P < 0.05) reduced Faith’s phylogenetic diversity on days 23, 25, 27, 29, 31, 35, and 39 in LBW piglets and days 23, 25, 27, 31, and 35 in NBW piglets. In terms of birth weight effect, NBW piglets showed significantly (P < 0.05) greater Faith’s phylogenetic diversity than LBW piglets on days 27, 29, 37, 39, and 41 in the antibiotics (AB) group, and days 27, and 41 in control (CON) group. Firmicutes_A was the most dominant phylum over the whole-time span of the experiment (Fig. 2C). Firmicutes_D and Proteobacteria were the subdominant phyla at birth and decreased in relative abundance over time. Relative abundances of Bacteroidota and Firmicutes_C increased from birth until the weaning period and from day 20 were the most dominant phyla next to Firmicutes_A. Similarly, *Escherichia_710834* (Proteobacteria phylum) was the most dominant genus at birth and decreased in relative abundance (Fig. 2D). *Prevotella* (Bacteroidota phylum) and *Megasphaera_*A_38685 (Firmicutes phylum), both Gram-negative organisms, increased in relative abundance over time and became dominant after weaning (Fig. 2D and E). Antibiotics significantly increased the relative abundance of *Parasutterella* and decreased the relative abundance of *Bifidobacterium*_388775 and *Ligilactobacillus* in at least two collection timepoints (ALDEx2 with q value of 0.05, Table S3). As expected, anaerobic bacteria were dominant in the gut microbial communities and there was a general increase in their relative abundance over time (Fig. 2F).

**FIG 2.**
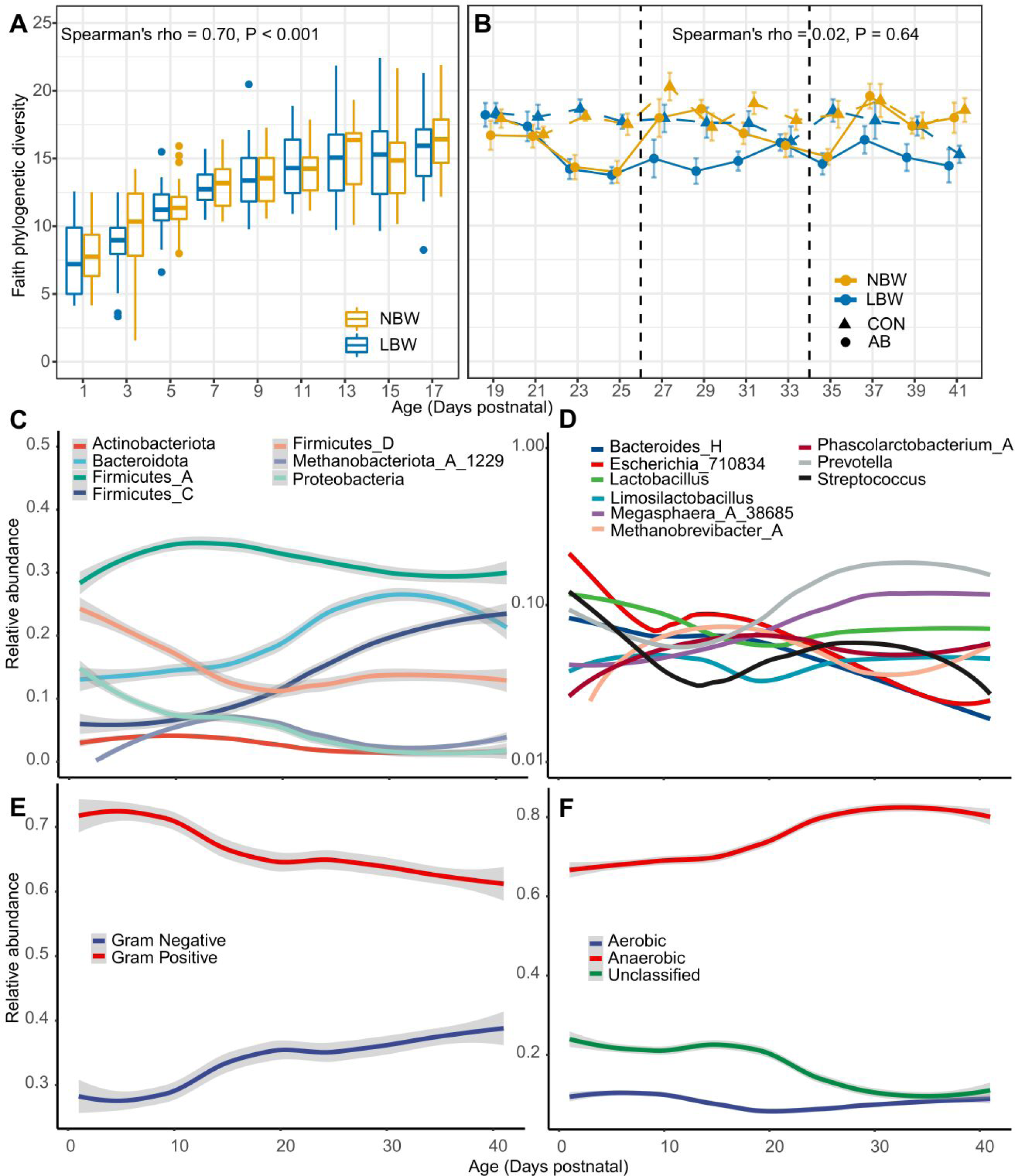
Descriptive overview of the swine fecal microbiome in the first 41 days of life. (A-B) Faith’s phylogenetic diversity of piglets in suckling stage (A) and weaning stage (B). (C-F) LOESS fits (95% CIs shaded in grey in C, E, and F) over time for the top seven phyla (C), the top nine genera (D), Gram-positive and Gram-negative bacteria (E), and aerobic and anaerobic bacteria (F).

### Antibiotics consistently change the gut microbiota structure

To visualize the changes in microbial community structure over time and to determine the effects of birth weight and post-weaning antibiotics on the swine fecal microbiota, we used Principal Component Analysis (PCoA) ordination plots in combination with Permutational Multivariable Analysis of Variance (PERMANOVA) on Bray-Curtis dissimilarity matrix (Fig. 3A). Age explained most of the variance in the gut microbiota succession in both suckling period (R^2^ = 0.10, P < 0.001) and weaning period (R^2^ = 0.10, P < 0.001), respectively. We then explored the longitudinal trends of beta diversity captured by the first two axes of independent PCoAs from all the samples (Fig. 3B). PCo1 and PCo2 tracked the development that microbial communities underwent during the early stages of piglets as evidenced by the progressive transition of early to late time points along the axes. Age-matched pairwise comparisons were conducted to explore the effects of birth weight and post-weaning antibiotics on the swine fecal microbiota at each time point. There were no significant effects of birth weight on the swine fecal microbiome during the suckling period (Fig. S2). From day 21 onward, the gut microbiota of piglets consuming diets containing antibiotics was statistically different (p < 0.05, R^2^ values between 0.07 and 0.11) from those on antibiotic-free diets on days 25, 27, 29, 31, 33, 35, 39, and 41 in LBW piglets, and days 21, 23, 25, 27, 29, 31, 33, 35, 37, 39, and 41 in NBW piglets (Fig. 3C). Birth weight had significant effects on swine gut microbiota on days 23, and 25 in CON group, and days 27, and 29 in AB group.

**FIG 3.**
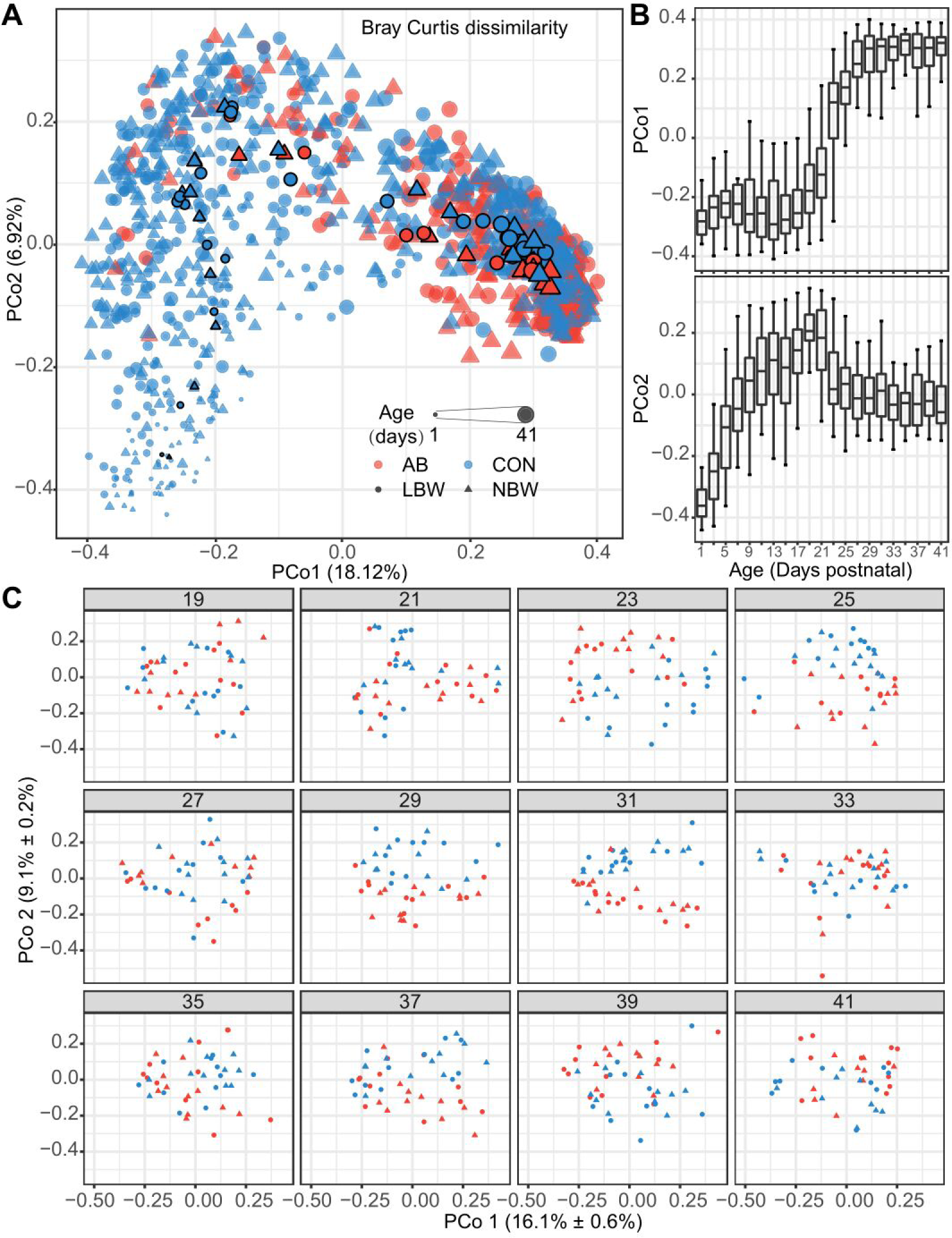
Birth weight and in-feed antibiotics are associated with swine fecal microbiome development. (A) Bray-Curtis dissimilarity principle coordinates analysis (PCoA) of all the samples collected during piglets growth. Point shape indicates birth weight, point color indicates post-weaning antibiotic use, and point size is proportional to age in days. Centroids were presented with closed lines. (B) Box plots shows PCo1 and PCo2 versus sample age for all piglets of all treatment groups. (C) Bray-Curtis dissimilarity principle coordinates analysis (PCoA) of samples collected during post-weaning stage faceted by age. Point shape indicates birth weight, and point color indicates antibiotic use.

### Individuality of the gut microbiota

Homogeneity of weaning piglets within each time point increased over time, as measured by a decrease in within-age Bray-Curtis dissimilarities (Fig. S3), indicating the stabilization of the microbiota development. Given that the relatively stable population composition and host-microbe interactions after 10 days succession post-weaning (37), we were able to quantify the intra- and inter-individual variation of the gut microbiota based on the last four observations of each piglet (Fig. 4). By analyzing overall gut microbiota composition using Bray-Curtis dissimilarity measured from the last four collection times, we observed more inter-individual variation than intra-individual variation (Fig. 4A), confirming that the gut microbiota is individualized. The individualized gut microbiota was also observed in the developmental phases, although both intra- and inter-individual variation decreased over time (Fig. S4). The degree of intra-individual variability was not affected by birth weight or antibiotics, possibly due to the homogeneous environmental, genetical, and nutritional backgrounds. However, we also observed that the degree of microbial variability over the last four times differed between the individuals (Fig. 4B). Individuals with the most intra-individual variability had a median intra-individual Bray-Curtis dissimilarity that was similar to the median inter-individual dissimilarity. To determine the intra- and inter-individual variance of individual microbial genera, we calculated the intraclass correlation coefficient (ICC) for the most abundant (with abundance > 0.1%, present in > 10% samples) genera (n = 72, Table S4). The total variance of abundance of each genera was negatively associated with the relative abundance of microbial genus (Spearman’s rho = -0.35, P = 0.003; Fig. 4C), indicating that the abundance of dominant genera was more stable. However, the median ICC was 0.35 (Table S4), indicating that there were more intra-individual variation of genus abundance than the inter-individual variation for the majority of the 72 most-prevalent genera. Among the most abundant genera with relative abundance > 1%, *Methanobrevibacter*_A and *Limosilactobacillus* had ICCs > 0.5 (Fig. 4D), indicating greater variability between individuals than within individuals, while *Mitsuokella*, *Alloprevotella*, *Megaspheara*_A_38685, *Lactobacillus*, *Cryptobacteroides*, *Gemmiger*_A_73129, *Blautia*_A_141781, *Phascolarctobacterium*_A, and *Prevotella* had higher intra-individual variations.

**FIG 4.**
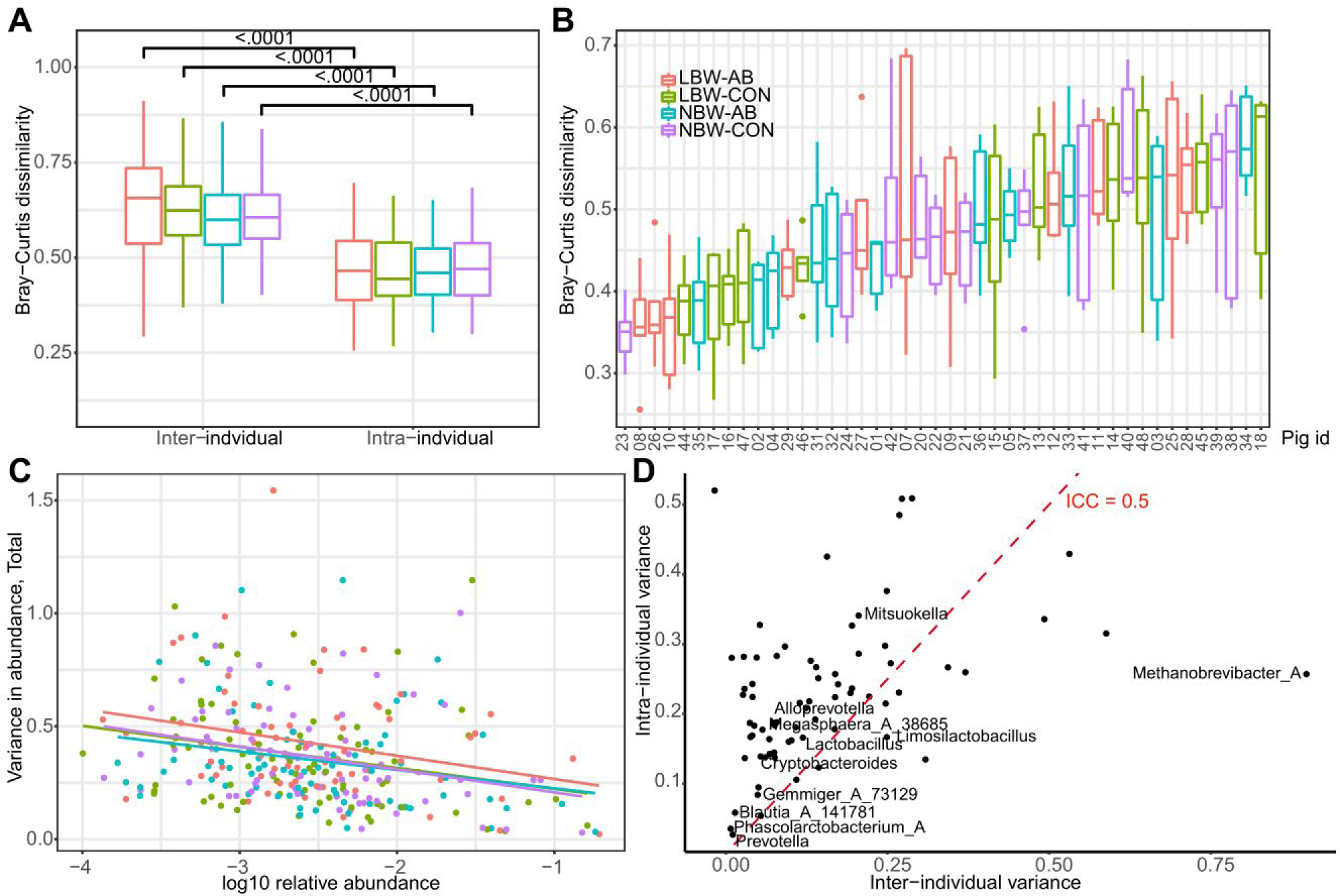
Patterns of inter- and intra-individual variation for microbial communities observed in the last four time points (days 35, 37, 39, and 41). (A) Boxplot of average intra- and inter-individual compositional variability using Bray-Curtis dissimilarity for piglets in different groups. (B) Compositional variability of the gut microbiota for the 44 individuals estimated by the Bray-Curtis dissimilarity. Boxplots are ordered by the median Bray-Curtis dissimilarity calculated for pairwise comparisons of the four samples from each individual (n = 6 values for each boxplot). (C) Total variance of individual microbial genera in last four collection times, plotted against the relative abundance. Dots are colored according to groups. (D) Relationship between the inter-individual and intra-individual components of total genus variance. Dashed line at intraclass correlation coefficients (ICC) 0.5 separate genus with high and low intra-individual variance, respectively. Dots indicate microbial genus.

### Persistent gut microbiota immaturity in LBW piglets

The maturation patterns of the gut microbiota were indicated using microbiota age predicted by the Random Forest regression models (Fig. 5A). We used a linear mixed-effects model including random intercepts and random slopes for individual piglets nested by their mothers, and evaluated birth weight, age, and antibiotic treatment as fixed factors and gender as a cofactor. There were no significant effects of birth weight on microbiota age during the suckling stage, but we found that microbiota age was affected significantly by birth weight (P = 0.03) after weaning. After running post-hoc pairwise comparisons, we found that within the CON diet groups LBW piglets had lower microbiota age than NBW piglets on days 23, 27, 33, and 41, and there were trending effects (0.05 < P < 0.1) on days 25, and 29 (Fig. 5B). Antibiotics reduced the microbiota age of LBW piglets on days 31 (P = 0.09), and 35 (P = 0.02), and reduced the microbiota age of NBW piglets on days 23 (P = 0.06), and 31 (P = 0.04) (Fig. S5).

**FIG 5.**
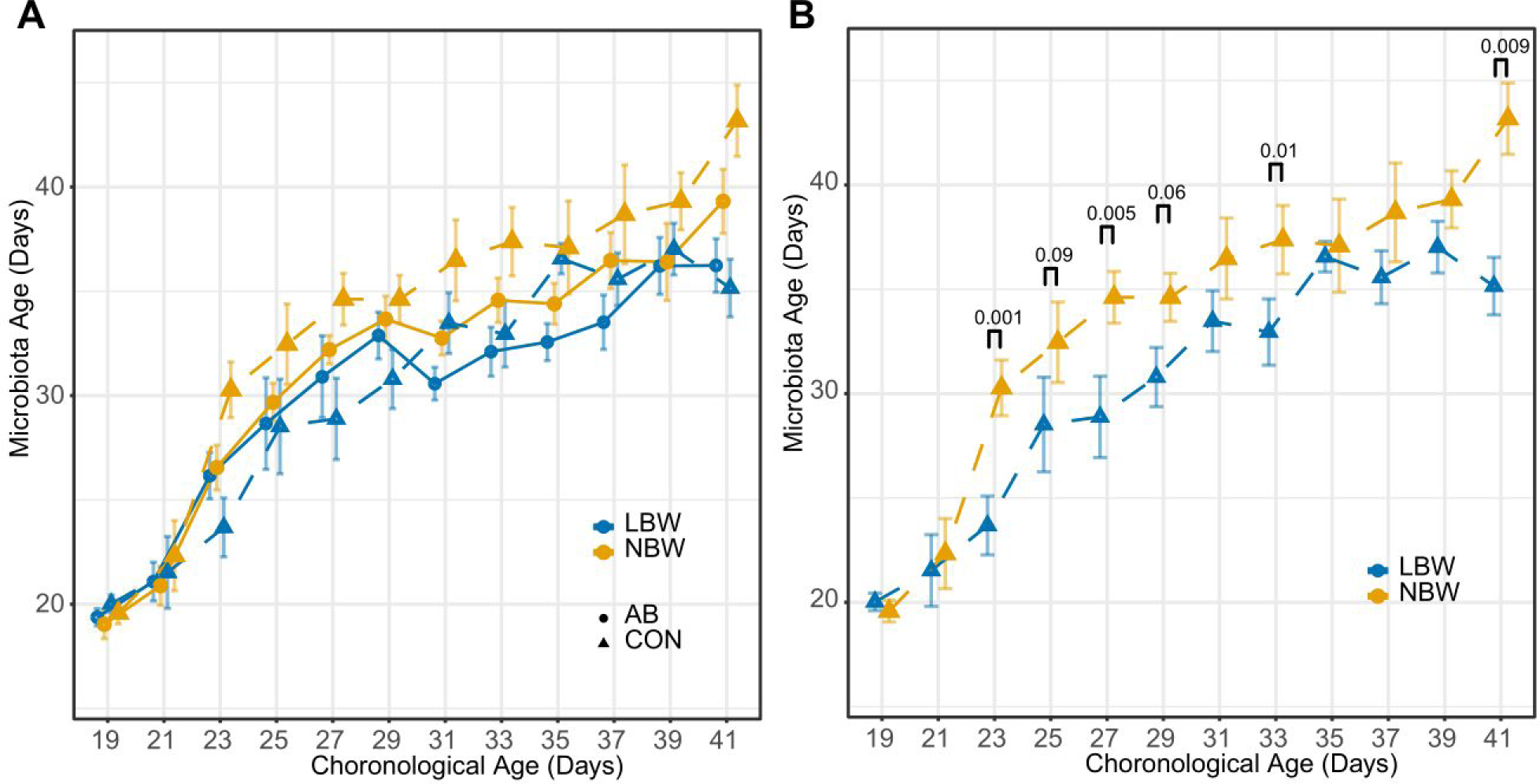
Maturation curve of swine fecal microbiota. Microbiota age predictions in all samples (A) and control samples (B) collected post-weaning. Point shape and line type indicate post-weaning antibiotic use, point color indicates birth-weight. Dashed lines with triangle points indicate control samples. Post-hoc P values are shown in panel (B). Data are means ± SEM.

### Fecal microbiota development follows an age-dependent pattern

Using the total cohort, we tested whether swine gut microbiota at different ages clustered according to age using Dirichlet Multinomial Mixtures (DMM) analysis, an unsupervised method to bin samples based on the structure of the microbial community. We identified 5 community clusters: 3 were common in samples from newborn piglets (clusters NB1, 2, 3), and 2 clusters were common in samples from post-weaning piglets (clusters PW1, 2). A clear age-dependent progression was observed when plotting the DMM clusters against the age of piglets (Fig. 6A). Next, we identified the five ASVs that contributed the most to each cluster and compared the relative abundance of these taxa within each cluster against all other clusters (Fig. 6B). As expected, a general increasing trend in alpha diversity was also observed in DMM clusters over the chronological order of their dominance (Fig. 6C). Consistent with the microbiota maturation analysis, LBW-CON piglets have less PW2 clusters (which we found is a more mature community type) during postweaning period (Chi-squared test, P = 0.03, Fig. S6A,B). All 5 clusters were observed in 40 piglets, two animals (#7 and #34) skipped NB3 and two animals (#9 and #25, both LBW-AB) never reached PW2 during the cluster transition over time (Fig. S6C). To quantify the transition of different clusters, we used a Markov chain-based approach to model the cluster transition probabilities (Fig. 6D). NB1 showed a high frequency of transition to other clusters, with the highest transition probability to NB2 (0.47), demonstrating the disappearance of NB1 after weaning. Although we observed a limited frequency of cluster transition from PW2 to PW1 (0.08), cluster PW2 had the highest level of self-transition, implying that cluster PW2 is more stable and mature compared to other clusters. The higher maturity and stability of PW2 was also indicated by an increase in PW1 samples in the last four timepoints of the animals with the most within-individual variability (Fig. S6D).

**FIG 6.**
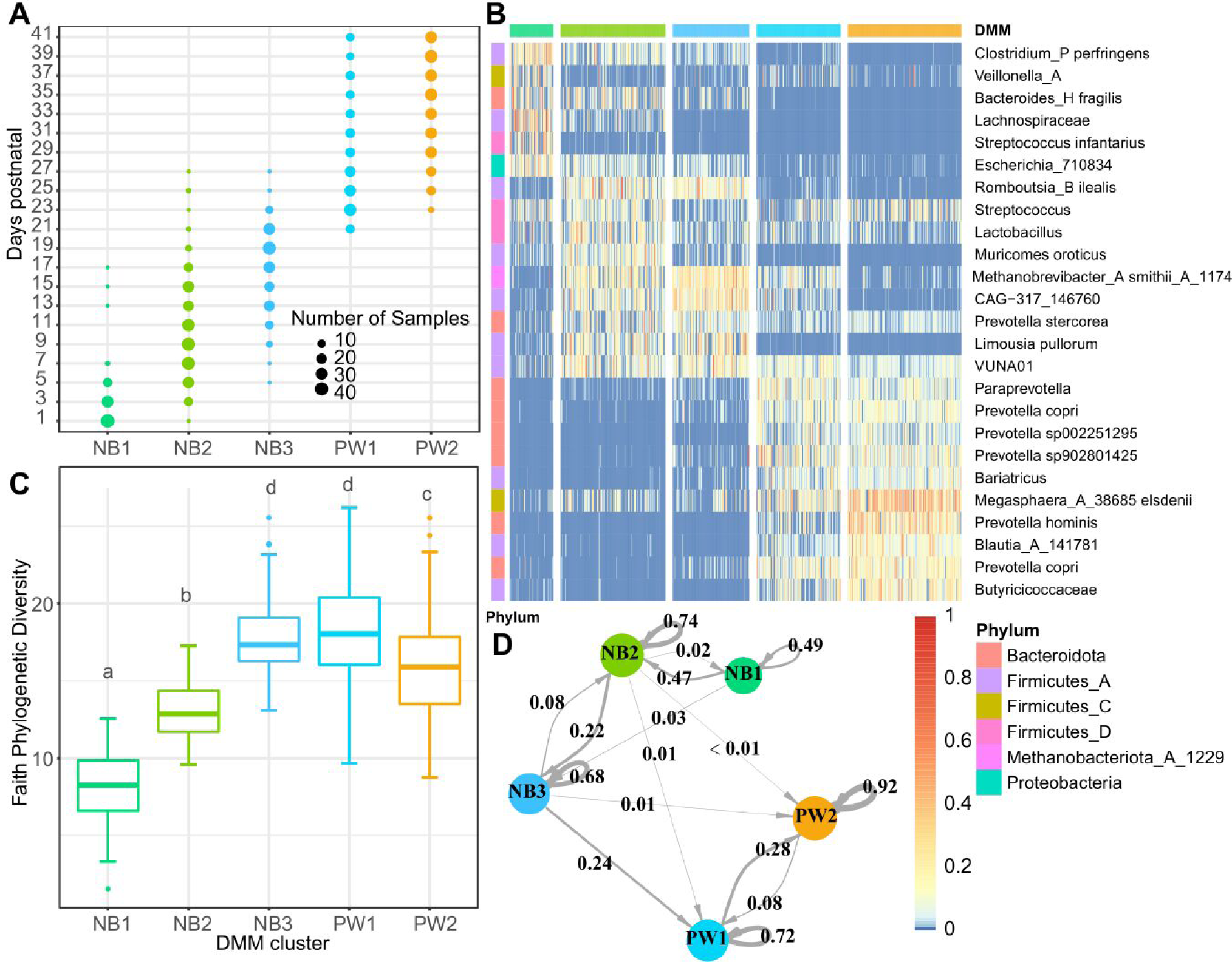
The gut microbial community development as measured by age-specific DMM clusters. (A) Distribution of samples in the identified 5 Dirichlet Multinomial Mixtures (DMM) clusters (x axis) using the DMM model at each time point (y axis). The size of each circle represents the number of samples within each cluster at each time point. (B) Heatmap showing the ASVs that contributed the most to each cluster (rows) and their relative abundance within each sample (columns). Samples are sorted by cluster assignment. (C) Box plots of Faith’s phylogenetic diversity within each DMM cluster, different lowercase letters indicate significantly different means (P < 0.05). (D) Markov chain with piglet-independent transition probabilities among clusters, in which arrow weights are proportional to the maximum likelihood estimate of the transition rate.

### Fecal microbiota assembly mainly shaped by deterministic processes

The neutral community model (NCM) was used to estimate the relationship between the occurrence frequency of ASVs and their relative abundance variations (Fig. 7A). This model can be used to infer the role played by stochastic processes in community assembly and quantify the importance of processes which are hard to observe directly but have large impacts on the microbial community. The NCM partially explained the community variance for LBW suckling piglets (R^2^ varied from -0.07 to 0.36, with the mean value of 0.22), and NBW suckling piglets (R^2^ varied from 0.03 to 0.32, with the mean value of 0.24). Less community variance was explained by NCM in weaning piglets, with R^2^ of 0.11 (-0.21 ∼ 0.28), 0.20 (-0.041 ∼ 0.33), 0.11 (-0.17 ∼ 0.26), and 0.16 (-0.01 ∼ 0.34) for LBW-CON, NBW-CON, LBW-AB, and NBW-AB weaning piglets, respectively. Despite the relative importance of stochastic processes increased over time within each growth period, the stochastic processes still explained limited variance in swine gut microbial communities assembly. Next, we used the infer community assembly mechanisms by phylogenetic bin-based null model analysis (iCAMP) framework to further partition the ecological processes into deterministic processes (heterogeneous selection, homogeneous selection, dispersal limitation, homogenizing dispersal), and stochastic processes (drift and others) (Fig. 7B). Our results support the prominent role of two deterministic processes, homogeneous selection and dispersal limitation, in driving the swine gut microbial community assembly. The mean relative importance of homogeneous selection in community assembly was 39.4% in suckling piglets, with a maximum of 60.4% in the NBW-CON group on day 1, and a minimum of 25.7% in the LBW-AB group on day 15. The mean relative importance of homogeneous selection in community assembly was 31.9% in weaning piglets, with a maximum of 42.2% in the LBW-CON group on day 41, and a minimum of 20.3% in the NBW-AB group on day 21. The mean relative importance of dispersal limitation in community assembly was 59.3% in suckling piglets, with a maximum of 72.9% the LBW-AB group on day 15, and a minimum of 36.8% in the NBW-AB group on day 1. The mean relative importance of dispersal limitation in community assembly was 67.3% in suckling piglets, with a maximum of 78.9% in the NBW-AB group on day 21, and a minimum of 56.7% in the NBW-CON group on day 33.

**FIG 7.**
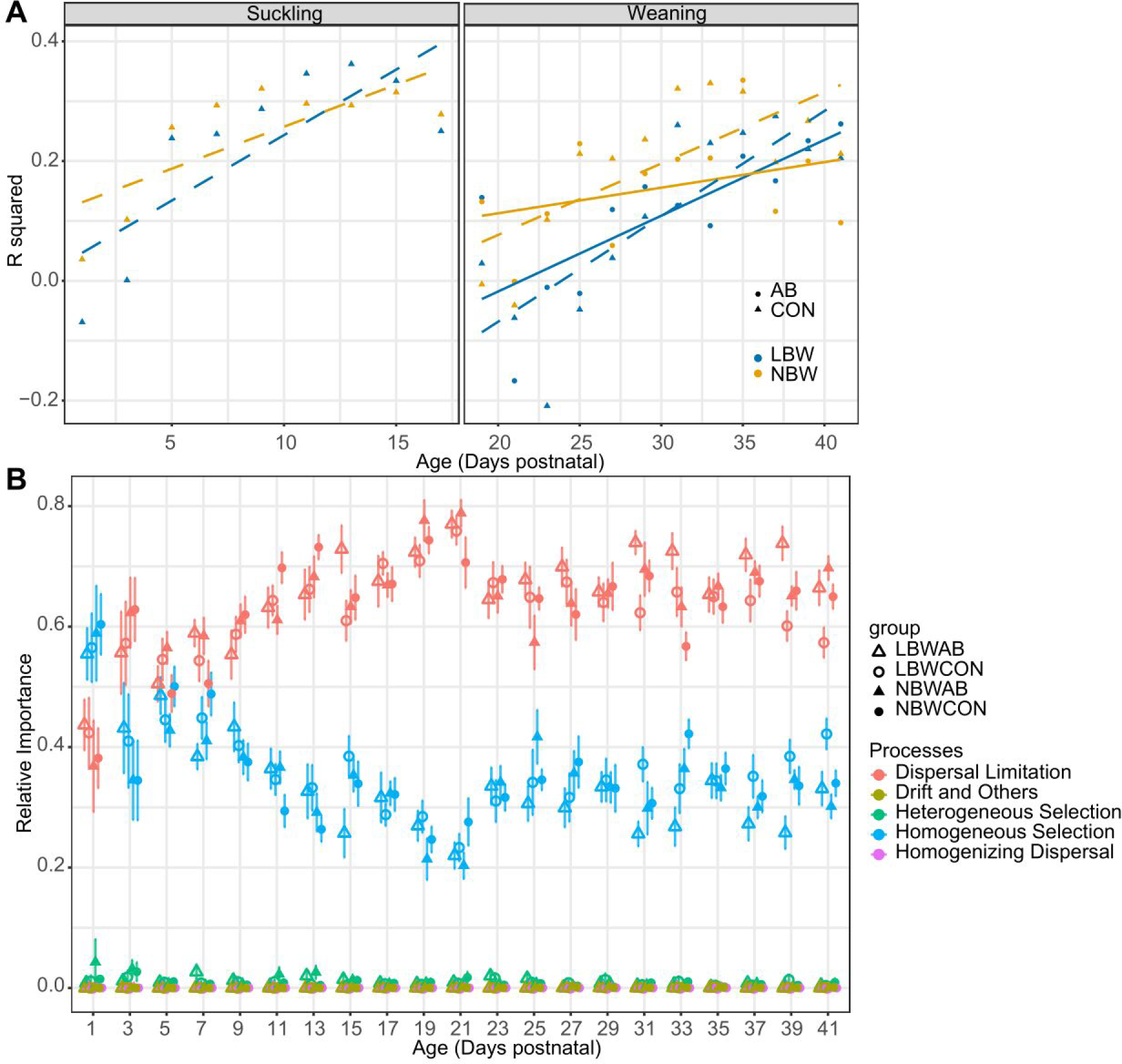
Ecological processes important to swine microbiome succession. (A) R squared indicates the fit of the neutral community model (NCM) of community assembly. Point shape and line type indicates antibiotics use, point and line color indicate birth weight. Dotted lines with triangle points indicate samples collected from piglets consuming diets without antibiotics. (B) Relative importance of different ecological processes in fecal microbial community assembly quantified by infer community assembly mechanisms by phylogenetic bin-based null model analysis (iCAMP) framework. Samples are grouped by groups at the end of the experiment. Point shape indicates sample groups, and point color indicates different ecological processes. Data are means ± SEM.

## DISCUSSION

Our observations provide additional understanding of the swine gut microbiome assembly and succession in the most dynamic periods of piglet growth. Repeated measurements, every two days, enabled us to investigate the temporal dynamics by capturing both individuality and generality of the gut microbiota (38, 39). We observed significantly negative correlations between age and the within-age Bray-Curtis dissimilarities in weaning piglets, indicating the gut microbiota is becoming more and more similar over the course of time (Fig. S3). Consistent with the current study, previous studies also indicated that the gut microbiota of piglets shifted quickly and reached relatively stable level 10 days after weaning (37). Stabilization of dietary composition and the development of the intestinal tract enables the succession of piglet gut microbiota to continue toward a stable and self-sustaining ecological community (23, 27).

Quantifying the intra- and inter-individual variation of the gut microbiota of each piglet has important practical implications. Individual differences are an essential consideration in animal experiments, despite providing a high degree of control over the genetic background and rearing environment (40, 41). We observed that the greater the intra-individual variation, the more unstable the gut microbiota (Fig. 4B, Fig. S6D). The gut microbial community of some piglets was highly variable suggesting that our capacity to detect individual differences or treatment effects may be diminished. Therefore, gut microbiome research requires large sample sizes to identify effects (42). Calculation of summary measurements from large sample sizes and longitudinal sampling could also be used to resolve the power issues (43, 44). Community succession continues in the gut microbial community even after it reaches a relatively stable state. Intrinsic dynamics indicate insufficient estimations of the equilibrium abundance from single measurements and highlight the importance of repeated measurements in microbiome research (38, 42). Moreover, community level descriptors such as beta diversity, enterotypes (DMM clusters here), or microbiota age are more stable than individual taxonomic biomarkers, and therefore may have better implications in treatment comparisons (33).

Along with intestinal development, comes the maturation of the gut microbiota, which is closely related to the healthy growth of the newborn (45, 46). In the current study, we used the random forest regression model trained by 12 previously published datasets (47) to quantify gut microbiota maturation. The independence training sets from our current experimental samples granted more confidence in the microbiota age predictions and comparisons. We observed that gut microbiota maturity (microbiota age) and growth performance (ADG) of piglets with low-birth weight piglets were significantly lower than those of normal piglets (Fig. 1, Fig. 5). Similarly, it was found previously, that early exposure to agricultural soil accelerated the early-life pig gut microbiota maturation and significantly increased ADG throughout the weaning period (48). The authors hypothesized that soil exposure enriched the plant-degrading genera and improved the fermentative potential which could explain the better growth performance (48). Moreover, piglets with more mature gut microbiota is associated with higher feed intake and reduced susceptibility to colonic inflammation (13, 45). However, the causality of the gut microbiota maturation and the growth performance in pigs needs to be further verified. The causal relationship of gut microbiota immaturity and childhood malnutrition was confirmed using gnotobiotic mice and pig models colonized with gut microbiota from healthy and malnourished children (35, 36, 49). Our results indicate that pig health and growth performance would likely be improved through gut microbiota development if the causality could be validated in the future.

There is a caveat that both our and previous observations were found in weaning piglets, and there was no correlation between microbiota maturation and growth performance in suckling piglets. This may be due to the fact that other factors, such as intra-utero growth retardation, milk competition, or gastrointestinal development have a much greater impact on growth performance than gut microbiota during the suckling stage (17–19). On the other hand, our current random forest model may be more generalizable in capturing the changes in gut microbiota in weaning piglets consuming plant-based diets, and less applicable for suckling piglets consuming milk (48). Moreover, when applying a random forest algorithm for the swine gut microbiota data, there are some limitations that still need to be resolved, such as feature selection when the microbial communities of different groups or across time points are highly similar (50).

Interestingly, antibiotics delayed the gut microbiota maturation yet increased the growth performance of weaning piglets in our study (Fig. S5). The above discrepancy to piglets consuming antibiotic-free diets emphasized the importance of determining the causal relationship between gut microbiota maturation and growth performance. This could also be explained that the promotion of growth performance by antibiotics and gut microbiota maturation may be through a different mechanisms of action (51). For example, we found that antibiotics significantly increased the relative abundance of *Parasutterella*, which is considered to be correlated with various health outcomes, such like succinate production, bile acid maintenance and cholesterol metabolism (52). However, *Parasutterella* made a negligible contribution to training the random forest model (47).

In addition to treatment-related effects, we observed an age-related pattern in the clusters, with maturation of the swine gut microbiome. The PW2 cluster showed its stability and maturity over other clusters based on the higher self-transition rate (Fig. 6D) and less intra-individual variability (Fig. S6D). In line with the microbiota age analysis, samples collected from LBW piglets eating CON diets showed fewer PW2 clusters, confirming the age-dependent stabilization pattern of the gut microbiota. Using neutral community and null models, we were able to quantify for the first time the ecological processes in swine gut microbial community assembly (Fig. 7). Homogeneous selection and dispersal limitation dominated the gut microbial community assembly over the entire suckling and weaning periods in this study, which are also observed to be dominant in human gut microbiome assembly (53). Homogeneous selection is due to a consistent selective environment among local scales, in this case the extremely similar intestinal environments and diet formulations. Dispersal limitation is due to limited exchange of microbes, indicating coprophagia played a very limited role in the transmission of gut microbes in suckling piglets and the successful isolation effect of single-cage feeding post-weaning (54). Deterministic process-driven community assembly patterns of the gut microbiota offer the possibility of successful interventions with community structure (30). The increasing proportions of stochastic processes over time indicates the importance of providing multiple interventions if attempting to direct gut microbial community composition.

In this study, we presented a comprehensive and quantitative analysis of the swine gut microbiota assembly and succession in 44 piglets sampled every other day. We observed individualized gut microbiota with substantial temporal variations, indicating the importance of large sample size and repeated measurements in swine gut microbiome research. We confirmed our hypothesis that the gut microbiota is immature in piglets with reduced birth weight and growth performance, which provides insights to intervene in animal growth through gut microbiota maturation. Swine gut microbiota assembly and succession are predominantly deterministic, with increasing stochastic processes over time. These findings provide a better understanding of microbial ecology and theoretical basis for practical applications. *[1167 words]*

## MATERIALS AND METHODS

### Animals and experimental design

All protocols used in the study were approved by the Purdue University Animal Care and Use Committee (West Lafayette, IN, USA). The pig experiments were carried out at the swine unit of the Animal Sciences Research and Education Center at Purdue University. A total of 44 piglets (22 low- and 22 normal-birth-weight piglets; LBW and NBW, respectively) were randomly selected for inclusion in the study based on farrowing date, litter, and sex. A NBW piglet was selected as a piglet with a birth weight within 1 standard deviation (SD) of the mean birth weight of the whole litter, whereas a piglet with a birth weight 2 SD below the mean was selected as LBW. Average birth weights for all LBW and NBW piglets in the current study were 1.19 ± 0.09 and 1.88 ± 0.20 kg, respectively. All piglets were weaned on day 18 postnatal. Following weaning, piglets were allotted to 4 groups based on a 2 × 2 factorial design according to birth weight (LBW and NBW) and post-weaning antibiotic treatment (AB for antibiotics, CON for control without antibiotics). The dosage of antibiotics was based on the recommended values for normal management of the Purdue swine farm. Neo-Terramycin^®^ 100/100 (neomycin-oxytetracycline) were added in the feed at therapeutic levels in the first two post-weaning weeks (850 ppm, and 550 ppm, respectively), and Mecadox^®^ 10 (carbadox) was added with a subtherapeutic level of 55 ppm during the third week of the weaning period (Table S1). Each group consisted of 11 replicates from 11 different sows, respectively. All piglets were caged in individual pens and fed the starter diet at the same nutritional level. All piglets were sampled using rectal swabs (PurFlock Ultra Sterile Ultrafine Flock Swab; Puritan, Guilford, ME) beginning at 1 day of age and repeated every two days until 41 days of age. All rectal swabs were immediately placed on ice, transported to the laboratory, and stored at -20°C until DNA extraction. Fecal scores (grade 1: hard, dry, and crumbly, and grade 5: watery diarrhea) were recorded for each piglet on each collection time and diarrhea piglet were classified if the fecal scores > 3 (55).

### DNA extraction and library preparation

Rectal swabs were processed to extract the bacterial and fecal content from the swab tip according to our previous publication (**56**). Briefly, the swab tip was mixed with 1 mL nuclease-free water using a vortex for 5 minutes, then centrifuged at 6000 × g for 10 minutes to form a pellet of the swab contents. The supernatant was then discarded, and the fecal pellet was stored at -20°C. DNA was extracted from fecal pellet with the QIAamp PowerFecal Pro DNA Kits (Qiagen, Hilden, Germany) according to the manufacturer’s protocol. Extracted DNA was quantified by NanoDrop and diluted to 10 ng/μL before library construction. The 16S rRNA gene sequence libraries using the Illumina MiSeq sequencing platform were constructed according to Kozich et al (57). Briefly, the V4 region of the bacterial 16S rRNA gene was amplified using 515F (GTGCCAGCMGCCGCGGTAA) and 806R (GGACTACHVGGGTWTCTAAT) primers. A mock community (20 Strain Even Mix 138 Genomic Material; ATCC^®^ MSA-1002^TM^) and water control were used in library construction. The amplified DNA, water, and mock community were normalized using SequalPrep Normalization Plate Kit (Invitrogen, Carlsbad, CA, USA) and pooled into equal volumes. The pooled samples were sequenced with Illumina MiSeq 2 × 250 bp paired-end sequencing at the Purdue Genomics Core Facility.

### 16S rRNA gene sequence analysis

Raw MiSeq fastq reads were imported into the QIIME2 (2021.11) for further analysis (58). Amplicon sequence variants (ASVs) were generated from demultiplexed paired-end reads with DADA2 (59). The forward sequences were trimmed at 13 and 250, and the reverse sequences were trimmed at 13 and 222. The mock community reflected the expected composition. A total of 12,438,593 high-quality reads from 6,290 ASVs were generated from 924 samples. After rarefaction of sample reads to 10,326, a total of 5,790 ASVs from 906 samples were included for downstream diversity analysis. Alpha and beta diversity analyses were conducted in QIIME2 using the q2-diversity plugin with core-metrics-phylogenetic command. Faith’s phylogenetic diversity was calculated to determine phylogenetic diversity in the microbial communities. The median Bray-Curtis dissimilarity calculated between samples from the last four collection times for an individual (i.e., 6 pair-wise dissimilarity values for the four samples collected from each individual) was used to quantify intra-individual compositional variability (34). The median Bray-Curtis dissimilarity obtained for the four samples from one individual versus all other samples from the same group was used to quantify inter-individual compositional variability (34). The taxonomy assignment of ASVs was based on the Naive Bayes classifier trained on the Greengenes2 database (2022.10) from 515F/806R region of sequences (60). The potential phenotypes of gram staining and oxygen tolerant of microbes in a community were predicted by BugBase (61). The maturation of the fecal microbiota was reflected by microbiota age calculated using predefined Random Forest regression models trained by publicly available data sets (47). To determine the key phases of microbiota progression, we used Dirichlet Multinomial Mixtures (DMM) clustering to bin samples based on the structure of the microbial community (62). The lowest Laplace approximation given to the negative log model evidence was used to estimate the optimal number of clusters (R package “DirichletMultinomial”, version 1.34.0). The intra- and inter-DMM cluster transition rates were quantified and visualized using a Markov chain by the method described previously (63). The quantitative framework Infer Community Assembly Mechanisms by Phylogenetic-bin-based Null Model (iCAMP) was used to examine the relative importance of different processes in the assembly of diverse microbe communities (64). The neutral community model (NCM), proposed by Sloan et al (65), was used in quantifying the importance of neutral processes in community assembly. The R^2^ parameter was used to represent the overall fit to the neutral model, all fitting statistics were calculated by bootstrapping with 1,000 replicates.

### Statistical analysis

All statistical analyses were performed in the R environment (4.2.1). General linear model (GLM) was used to analyze the effects of birth weight and post-weaning antibiotics on average daily gain (ADG). Significant differences in Faith’s phylogenetic diversity and microbiota age were calculated using linear mixed-effects model with the ‘mixed’ function of the ‘afex’ package. Differentially abundant microbial taxa were identified using ANOVA-Like Differential Expression (ALDEx) analysis using the q2-aldex2 plugin in QIIME2 (66). The effects of birth weight and post-weaning antibiotics on the microbial community structure were assessed using the Bray-Curtis dissimilarities with permutational multivariable analysis of variance (PERMANOVA) using ‘adonis’ function of the ‘vegan’ package. The R package ‘pairwiseAdonis’ was used to compare the Bray-Curtis dissimilarities within each sampling time. Comparisons between intra- and inter-compositional variance were analyzed using GLM. Mean and variance for each genus abundance were modelled by partitioning the total variance into inter-individual variance and intra-individual variance. Intraclass correlation coefficient (ICC), defined as the ratio between inter-individual and total genus abundance variance, was calculated using a one-way ANOVA fixed effects model based on log10 transformed relative abundance, as applied in the function ‘ICCest’ of the ‘ICC’ package. We ran the Shapiro-Wilk test for normality analysis. The Kruskal-Wallis test was used to assess changes in alpha diversity between DMM clusters, followed by a multiple comparison using the Wilcoxon rank-sum test with Benjamini-Hochberg (BH) p-value adjustment due to multiple testing. The chi-square test of independence was used to determine if there is a significant association between groups and DMM cluster distributions. Linear models were performed using the ‘lm’ function, Spearman’s rho were calculated using the ‘cor.test’ function from the “stats” package. The results were considered significant when P values were **≤** 0.05.

## Data availability

All 16 rRNA gene sequencing data are available at the NCBI Sequence Read Archive under BioProject No. PRJNA993266.

## SUPPLEMENTAL MATERIAL

Supplemental material is available online only

**FIG S1-S6**, PDF file, 1.84 MB.

**TABLE S1-S4**, XLSX file, 33.7 KB

## ACKNOWLEDGEMENTS

This project was supported by funding from Purdue University to T.A.J.

WD and TAJ conceived the study, WD, PO, RECM, TS, and TAJ collected the swab samples, WD performed the data analysis, WD, BR, and TAJ discussed the results, WD wrote the manuscript, TAJ edited the manuscript. All authors read and approved the final manuscript.

The authors declare that they have no competing interests.

## FIGURE LEGENDS

**FIG S1** Diarrhea incidence of animals on each day. Point and line colors indicate birth-weight. Line type indicates the use of antibiotics. Dashed lines indicate control samples. **FIG S2** Bray-Curtis dissimilarity principle coordinates analysis (PCoA) of samples collected during suckling stage faceted by age. Point color indicates birth weight.

**FIG S3** Within time Bray-Curtis dissimilarities of piglets in suckling stage (A) and weaning stage (B). LBW, low-birth-weight; NBW, normal-birth-weight; AB, antibiotics; CON, control. Pearson’s coefficients (r) were based on the linear correlation between within group distances and age.

**FIG S4** Boxplot of average intra- and inter-individual compositional variability using Bray-Curtis dissimilarity for piglets in different groups in days 19, 21, 23, and 25 (A) and in days 27, 29, 31, and 33 (B). (C, D) Patterns of the change of inter- and intra-individual variation over time.

**FIG S5** Effects of antibiotics on the maturation curve of swine fecal microbiota. Microbiota age predictions in LBW samples (A) and NBW samples (B) collected post-weaning. Dashed and solid lines indicate control and antibiotic-treated samples, respectively. Post-hoc P values are shown above each age-matched comparisons. Data are means ± SEM.

**FIG S6** The number of samples for each Dirichlet Multinomial Mixtures (DMM) cluster appeared in suckling (A) and weaning (B) stages, respectively. LBWCON piglets have less PW2 clusters during postweaning period (Chi-squared test, P = 0.03). (C) DMM clusters for each individual at the 21 visit times. Samples are colored according to the assigned community cluster. Dashed lines between days 17 and 19 indicate weaning age (day 18). (D) DMM clusters at the last four time points for each individual ordered by the increasing intra-individual compositional variability. LBW, low birth weight; NBW, normal birth weight; AB, antibiotics; CON, control. NB1-3, new born; PW1-2, post-weaning.

